# Combining deep learning and automated feature extraction to analyze minirhizotron images: development and validation of a new pipeline

**DOI:** 10.1101/2021.12.01.470811

**Authors:** Felix M. Bauer, Lena Lärm, Shehan Morandage, Guillaume Lobet, Jan Vanderborght, Harry Vereecken, Andrea Schnepf

**Affiliations:** Institute of Bio-and Geosciences: Agrosphere (IBG-3), Forschungszentrum Jülich GmbH, 52425 Jülich, Germany; Institut für Bodenkunde und Standortslehre, Universität Hohenheim, 70559 Stuttgart, Germany

**Keywords:** high-throughput phenotyping, minirhizotron, roots, workflow

## Abstract

Root systems of crops play a significant role in agro-ecosystems. The root system is essential for water and nutrient uptake, plant stability, symbiosis with microbes and a good soil structure. Minirhizotrons, consisting of transparent tubes that create windows into the soil, have shown to be effective to non-invasively investigate the root system. Root traits, like root length observed around the tubes of minirhizotron, can therefore be obtained throughout the crop growing season. Analyzing datasets from minirhizotrons using common manual annotation methods, with conventional software tools, are time consuming and labor intensive. Therefore, an objective method for high throughput image analysis that provides data for field root-phenotyping is necessary. In this study we developed a pipeline combining state-of-the-art software tools, using deep neural networks and automated feature extraction. This pipeline consists of two major components and was applied to large root image datasets from minirhizotrons. First, a segmentation by a neural network model, trained with a small image sample is performed. Training and segmentation are done using “Root-Painter”. Then, an automated feature extraction from the segments is carried out by “RhizoVision Explorer”. To validate the results of our automated analysis pipeline, a comparison of root length between manually annotated and automatically processed data was realized with more than 58,000 images. Mainly the results show a high correlation (*R*=0.81) between manually and automatically determined root lengths. With respect to the processing time, our new pipeline outperforms manual annotation by 98.1 - 99.6 %. Our pipeline,combining state-of-the-art software tools, significantly reduces the processing time for minirhizotron images. Thus, image analysis is no longer the bottle-neck in high-throughput phenotyping approaches.

## Introduction

Roots are an essential component of the global biosphere. They are mainly responsible for the acquisition of the resources water and nutrients for the entire plant. In most ecosystems, these resources are the limiting factors for growth of plant organs and yield (1). Water and nutrient uptake are directly linked to the parameters defining the root system, like length, diameter or branching. Therefore, collecting information about the root system becomes increasingly significant. In order, to improve water and nutrient uptake of plants for specific soil and climatic conditions, it is essential to obtain information about the root system architecture of plant species that have been shown to be beneficial for the given conditions (2). For plant breeding, this will help to develop new genotypes which are able to cope better with e.g. drought-stress and are more efficient in nutrient uptake (3). This will not only help to increase the arable area for certain species, it might also lead to higher yields. Especially this applies to locations with less suitable environments for a highly productive agriculture. The negative impact on the soil should be minimized at the same time (4).

The direct observation of roots is difficult, because the root system is surrounded by soil, making it impossible to visually measure the roots. To avoid that measurements heavily disturb the plant and its environment, preparations planned in advance, like the installations of rhizotubes, or the the construction of a minirhizotron, are crucial. (5). Minirhizotrons are useful tools to collect data about the root system without disturbing the environment of the roots or the plant itself. Moreover, they allow root observations over the whole vegetation period at a high temporal resolution and the comparison of different vegetation periods and crop types. Transparent rhizotubes, installed below ground, function as a window in the soil. Guided camera-systems provide photographs of the roots and the surrounding soil. Consequently, the non-invasive root measurements can be repeated multiple times during the growing period under *in situ* conditions. However, large minirhizotron facilities include tubes in different depth-levels. Measurements in several depths and time lapse observations result in big datasets that often consist out of 10,000 images and more (6). Images provided by minirhizotrons strongly differ from e.g. root scans gained from excavated and washed roots (7). Various soil conditions around the tubes in different depths lead to a wide range of heterogeneous images with different characteristics. Beside the actual roots, soil structures and disturbing fragments, including small animals, are depicted. Different soil conditions in various depths and at varying locations lead to varying color and light conditions and therefore make the automated processing of minirhizotron-images a challenging task (8).

To analyze roots mainly two steps are needed, the thresholding of root objects and the object quantification (9). Due to the heterogeneity within minirhizotron images, the thresholding is very complicated. Different analysis approaches emerged, represented by a numerous collection of software tools, designed to extract the information about the root system (10). These tools work manually, semi-automated or automated. Manual annotation tools for minirhizotron images, like “WinRhizoTRON” (Regent Instruments Incl.), rely on the human interaction with each individual image taken, to track each root by hand. It requires the user to follow every root depicted in the image by hand and mark start, branch and endpoints. Semi-automated and automated approaches with software-tools exists to facilitate and speed-up the post-processing of the images (8). Filter algorithms used to increase the contrast between root and background and to find root structures by typical geometrical shapes, were proposed by several authors (7, 11, 12). Semi-automated software like “RootSnap!” (CID Bioscience) and “Rootfly” (13) require a manual annotation, but also provide root suggestions by a filter created on an initial dataset. Consequently, most of these programs are strictly limited to certain type of images, like high-contrast root scans (14). Eventually, this has the consequence that the annotation of the roots in most minirhizotron images needs to be done almost exclusively manually. Depending on the number of images taken and the number and length of roots, the manual and semi-automated analysis can take weeks or even months. Previous studies found that the estimated amount of minirhizotron images, annotated with an annotation software, was between 17 and 38 images h^−1^ (15). Adapted to the working routine with “Rootfly”, it takes 1-1.5 h annotation time for an image area of 100 cm^2^ depicted soil (16). Further, the results underlie the subjectivity of the annotator.

Deep learning has developed to the Gold Standard of machine learning methods within the recent years. Deep neural networks are able to learn from big datasets and provide outstanding results on complex cognitive challenges, even beating human performance in some application fields (17). Convolutional Neural Networks (CNN), a subclass of deep learning, are created to deal with data in the shape of multiple arrays and are therefore suitable for high-dimensional data like images (18). They have the potential to perform a decent automated detection of a certain structure within a heterogeneous and noisy dataset (19). Transferred to the analysis of minirhizotron images, CNNs should have the capability to precisely identify and segment roots in images, where the roots cannot be segmented sufficiently by e.g. explicit programmed thresholds or filter algorithms. CNNs were already used successfully to localize plant organs, including roots (20–23). However, the use of CNNs has mainly been proved on data originating from controlled environment, like lab experiments (14). Furthermore, they are often limited to the use of one or a few fixed pre-trained neural network models (24), or they are not easily usable for non IT-professionals (25). The main reason for this is the required knowledge and competences in machine learning and programming needed to create a CNN-based system. Especially the data partition between training and validation, the process of annotation and the setup of network architecture make the use of CNNs complicated (26). Although the use of CNN is promising for root segmentation, it is not subject of many published studies and not yet widely used as phenotyping tool for root traits. To make the advantages of CNNs widely utilizable, a software, combining the annotation, training and segmentation process with CNN together in an interface easy to handle, is the key for general use of neural networks for automated root segmentation. The recently published software tool “Root-Painter” is one of the most promising approaches for this task (27).

However, fast and reliable segmentation is only the first step of root analysis. For the root quantification another tool is required to obtain morphological and topological features from segmented images. For this task conventional automated root analysis tools, like “WinRhizo” (Regent Instruments Incl.) and “IJ_Rhizo” (28) can be used. Recent progress in the development of root-system feature extraction from high-contrast images or scans have resulted in new software tool with the ability of extracting multiple features with a high precision. On the front line of current developments is the new software “RhizoVision Explorer”, providing the functions to accurate skeletonization a high-contrast segmented image, to correct the skeleton and deriving several features from it (29).

The aim of our study is to develop a generally applicable, automated analysis pipeline, based on state-of-the-art technologies and software to extract root traits from minirhizotron images. This includes data annotation for neural network training, segmentation and feature extraction. The automated analysis pipeline has to meet the requirements in *i*) availability and feasibility, ii) accuracy and comparability, iii) speed and efficiency. It was an important requirement to us that this workflow should be feasible for root scientists, who only have basic knowledge in programming or computer science. This workflow should make fast root phenotyping easily accessible for newcomers in root science and lower the time and effort needed to get into the topic. Therefore, it relies on already published software. This workflow further should underline the practicability of deep learning phenotyping tools for the scientific root analysis routine. All software required to use this automated root image analysis pipeline are freely available and easy to operate. Another key advantage of our study is the scope of data used for validation and comparison and the concomitant claim to a general validity of this pipeline. To test and validate the automated analysis pipeline, datasets obtained from several years and two minirhizotron facilities were processed and compared to previously manual annotated data (6, 30–32). Previous studies evaluating the results of a CNN-automated image analysis for root images originating from (mini)rhizotrons used between 40 - 857 images (16, 24, 33). In our test we evaluated the results of more than 131,000 images of which we used more than 58,000 for a direct one-to-one comparison of manual human annotation to our automated analysis pipeline. The images represent different *in situ* conditions. In this paper we will present the detailed procedure on operating the automated analysis pipeline and compare its performance to a previously done manual annotation for a decent evaluation.

## Methods

### Experimental test site

The data used for the automated analysis pipeline were collected at the two minirhizotron facilities at the Selhausen test site of the Forschungszentrum Jülich GmbH (50°52’07.8”N, 6°26’59.7”E), Germany (34, 35). The field, in which the minirhizotron facilities are located, has a slight incline with a slope of under 4°. The two minirhizotron-facilities are approximately 150 m apart. The minihizotron facility located at the top of the field is hereafter referred to as RUT (rhizotron upper terrace) and the minirhizotron at the lower part of the field as RLT (rhizotron lower terrace). The thickness of the soil layer with silty loam texture varies strongly along the field-slope. While it is not present at the top, its thickness at the bottom is up to 3 m. At R_UT_ the gravel content is 60 % while at R_LT_ it is only 4 %. Both facility contain 54 horizontally installed, transparent tubes with each a length of 7 m and an outer diameter of 6.4 cm. The tubes are separated into three plots with each three vertical, slightly shifted (10 cm) rows of six tubes, where three different treatments can be studied. The tubes in each row are installed in −10 cm, −20 cm, −40 cm, −60 cm, −80 cm and −120 cm depth. Past treatments include different irrigation patterns (sheltered, rainfed, irrigated), different sowing densities and dates (later sowing in sheltered plot), or cultivar mixtures (two single cultivar treatments and one mixture). The two minirhizotron facilities were installed in 2012 (RUT) and 2014 (RLT), respectively. Further construction details are explained in (6).

### Data acquisition

Two different camera systems manufactured by Bartz (Bartz Technology Corporation) and VSI (Vienna Scientific Instruments GmbH) were used to capture the root images in the minirhizotrons. Both camera-systems are designed to be used manually. A regular measurement produces 40 images per tube. 20 images are taken 80° clockwise and 20 images 80° counter-clockwise from the tubes top point (6, 30, 36, 37). In this study, the collected images of three crop growing seasons from 2015/16, 2017 and 2018 were taken into account. Depending on the year and measurement date either the Bartz- or the VSI-system was used. The crops cultivated at the test site and used for this study were *Triticum aestivum* cv. Ambello in 2015/16 (winter wheat) and in 2017 and 2018 *Zea mays* cv. Zoey. Table 1 gives an overview on camera system used, the resolution of the images, measurement years, measured time period and cultivars observed. Depending on crop growing season, the total amount of measurement dates varied between 17 and 38. The amount of images, taken at one measurement date, varied according to the amount of tubes measured at this measurement date (Table 3). This was depending on the state of vegetation evaluated in field.

**Table 1.** Overview of the camera-systems and experiment timeline of minirhizotron images acquisition

| camera system | Bartz | VSI |
| --- | --- | --- |
| original resolution (px) | 754 x 510 | 3280 x 2464 |
| converted resolution (px) | 1508 x 1020 | 2060 x 2060 |
| real size (mm) | 16.5 x 23.5 | 20 x 20 |
| growing season | 2015/16 & 2017 | 2017 & 2018 |
| culture | 2015/16: *Triticum aestivum* cv. Ambello<br>2017: *Zea mays* cv. Zoey | *Zea mays* cv. Zoey |
| time period | 16/11/15 - 23/06/16<br>23/06/17 - 12/09/17 | 08/06/17 - 22/06/17<br>23/05/18 - 23/08/18 |

Over the past years, the root images collected in the minirhizotron facilities in Selhausen were analyzed manually, using “Rootfly” as a semi-automated tracking tool for the root length and root counts (6, 13, 30, 31, 37). In this study the images of the years 2015/16, 2017 and 2018 were analyzed. The manual annotation of 2015/16 and 2017 has been already published in (30, 32). Further a sub-sample of the root images was manually annotated by two persons separately. 1,760 images were used for the comparison between both annotators, and the annotators and the results of the automated analysis pipeline.

### Software tools

Our proposed automated minirhizotron image analysis pipeline is based on two software tools for the segmentation (27) and the automated feature extraction (29). Furthermore, scripts to convert the segmented images and analyze the outcome are available. For a easy accessibility all scripts are available together within the GUI of the executable “RootAnalysisAssistance”.

#### Segmentation

“RootPainter”, a software tool for the deep learning segmentation of biological images with an included annotation function provides an interactive training method within a GUI, using a U-net based CNN. U-net was developed to train with less images for a more precise segmentation and is therefore suitable when it comes to images where the manual annotation is especially time and labor consuming (16, 38). “RootPainter” was developed to make data-creation, annotation and network-training accessible for ordinary users. It provides a dataset creation function, which allows an easy selection of training images and cropping them in multiple tiles and to a suitable size for the interactive training. The training mode, provides an interactive graphical platform to manually annotate a small part of the dataset and create a neural network model. Further, a mode to segment whole image directories at once is provided. For training and segmentation a Graphics Processing Unit (GPU) is required (27). However, a full minirhizotron image analysis is based on two main components, the segmentation and the root trait extraction. Although “RootPainter” provides an inbuilt function for basic root trait extraction based on the previous segmented images, it does not provide e.g. a skeleton correction function and a comprehensive feature extraction including multiple root traits. For our pipeline the feature extraction part should provide multiple morphological and topological root features with a high accuracy. Furthermore, the possibility of a systematic correction function should be implied. Therefore a platform fulfilling these requirements was used for feature extraction.

#### Feature extraction

“RhizoVision Explorer” represents the current state-of-the-art technology with a sophisticated automated root traits extraction from segmented root images, by combining the abilities of several existing root image analysis platforms. This includes skeletonization of the segments, filter, filling, smoothing and pruning functions (29, 39). However, like most programs for automated root system analysis it is built for the use with binary images or high contrast scans and therefore not suitable for minirhizotron images. The capability of “RhizoVision Explorer” are nevertheless useful when applied to already segmented minirhizotron images.

### Analysis pipeline

The starting point for the automated analysis pipeline is a directory containing the raw images captured at the minirhizotron facility. The pipeline was run on a GPU-server with 4 Nvidia GeForce RTX 2080 Ti (NVIDIA Corporation). As client, a computer with an Intel i5-8265U processor and 24GB RAM, operated on Windows 10, was used. However, it is also possible to run the pipeline on one machine, if there is a GPU with CUDA available. An overview of all following steps is explained in Figure 1.

**Fig. 1.**
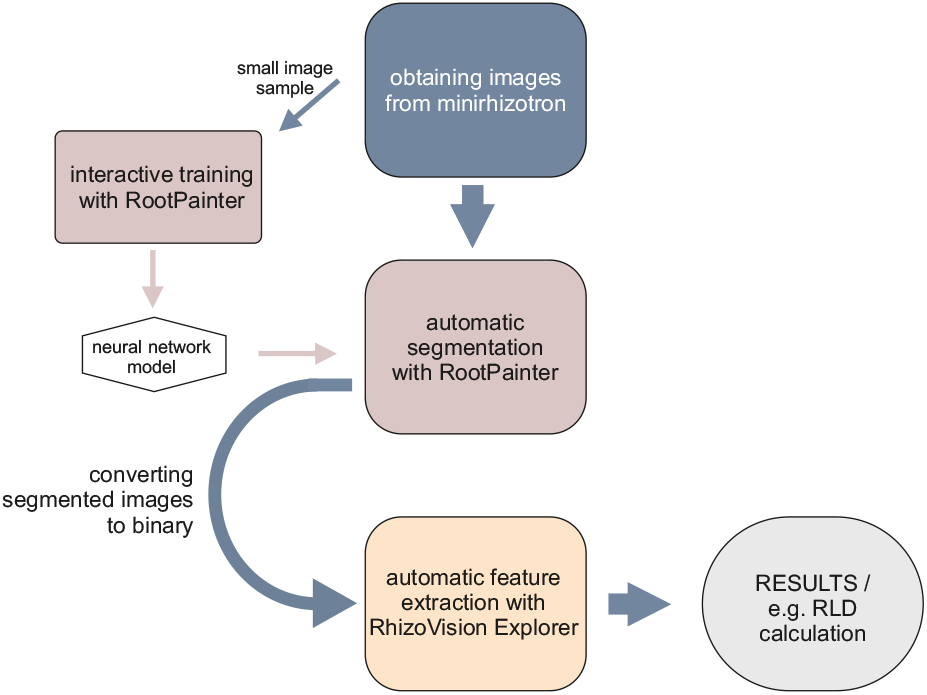
Schematic overview of the workflow of the automated analysis pipeline starting with image acquisition in the minirhizotron facility.

### Pre-processing

The first step of the pipeline is the preprocessing of the images. Depending on the camera-system either an up- or down-scaling and a distortion correction is performed. In the same step a labeling and sorting of the images is done automatically. If the images are already ready to use, this step can be omitted.

### Training

This step is only needed if no suitable neural network model exists for the targeted dataset. The process, to train a model for root segmentation starts with the creation of a training dataset and subsequently a new project in “Root-Painter”. We highly recommend to balance the training data according to the factors influencing image visually. In our case the training dataset for one model contains a balanced amount of images from two different minirhizotron facilities, respectively soil types, depths, tubes and dates. We used only a small amount of images from all available images. For each camera system a separate model was trained. The annotation can be done in the GUI. The roots are annotated as “foreground”, soil and other not root-belonging fragments as “background”. After the training is started, “RootPainter” automatically creates neural network models, depending on the annotation done previously. The progress can be seen in real time, because “RootPainter” provides previews of the segmentation done by the actual model. These proposals can be corrected and supplemented by the user. The training procedure used in this study is the “corrective training”. It is intended for large datasets and therefore suitable for the minirhizotron image data. Essentially this training approach starts with annotating a few images in detail and then continue with correcting only the false-positive and false-negative suggestions of the current model. Further details and instructions are explained in (27).

### Segmentation

The fully-automated segmentation is done with a model previously trained with a small selection of images from the corresponding measurements. To perform the fully-automatic segmentation, all images have to be located in one directory. The segmentation process itself is started from the “RootPainter” main menu. The result is a root segment (Figure 2.1,Figure 2.2).

**Fig. 2.**
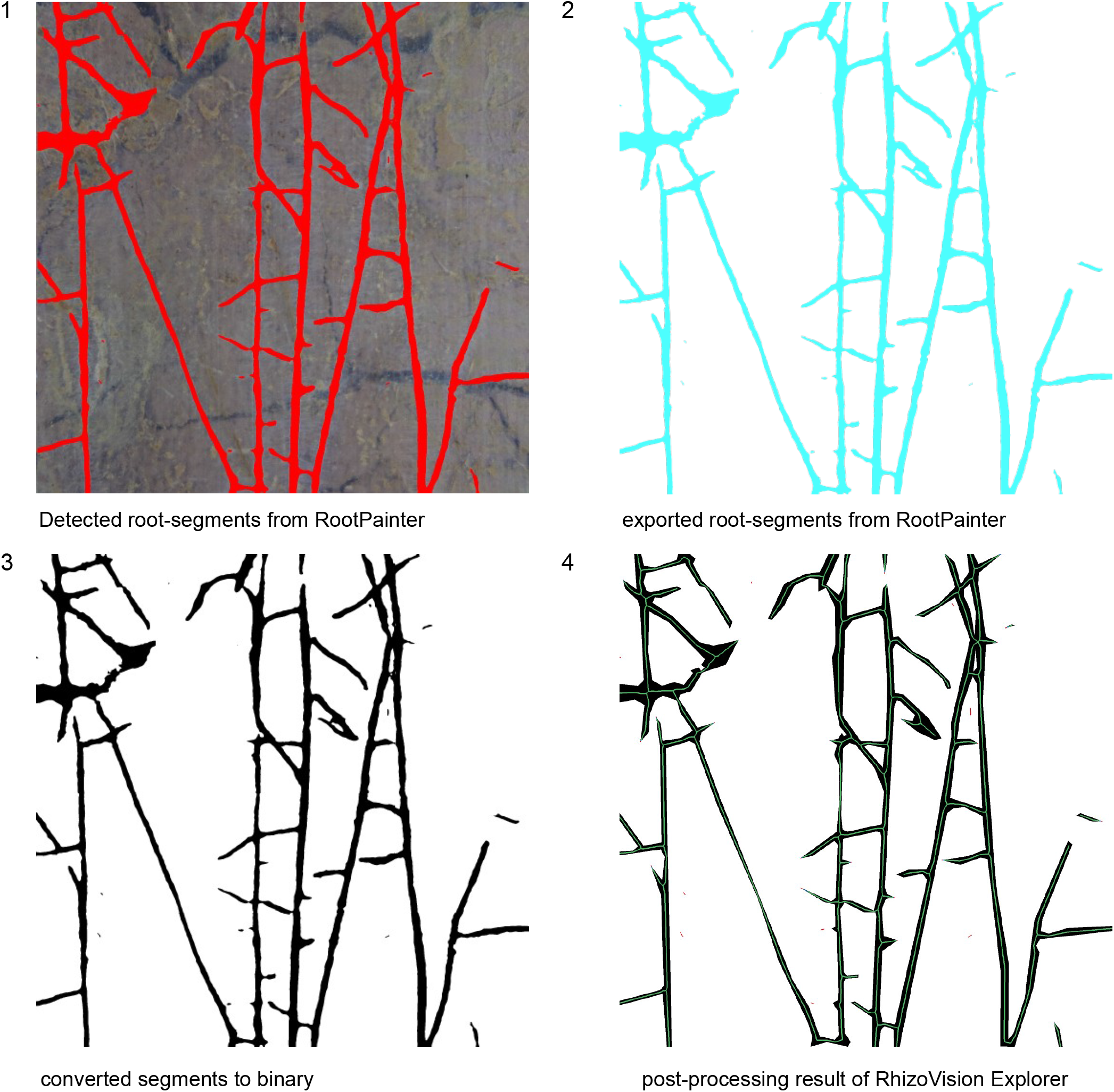
Example for one image processed by the automated root analysis pipeline. (1) The roots are “detected” by RootPainter according to the previous trained model. (2) The segmented image is exported and (3) converted to binary. (4) The last step is the skeletonization and feature extraction with RhizoVision Explorer.

### Converting

To import the segmented images into “RhizoVision Explorer” in the next step, it is essential to convert the images to binary, otherwise the images are not loaded properly (Figure 2.3). This step is performed by a conversion-script, which converts the mono-colored segmented images to black and white images and reduces the images information to binary by only giving information for either black or white pixels. The conversion-script is available as python script or within the *RootAnalysisAssistance-GUI*. It is possible to either browse the image-folders to convert manually, or to process the conversion of a certain image directory in a batch mode. This option is suitable for fast processing a huge amount of segmented images.

### Feature extraction

The final step is the feature extraction, performed by “RhizoVision Explorer”. This is also done in batch mode. The threshold of the non-root filter, hole filling, edge smoothing and pruning was chosen in a standardized way and uniform for each parameter, depending on the resolution of the image. For the images resulting from the Bartz-system the threshold is 13 px and for the VSI-system 20 px. This results in filtering parts smaller then 0.2 mm and filling holes bigger than 0.2 mm. To minimize the influence of segmentation mistakes at the boarder between root and soil and thus reduce the false detection of non-existent laterals, the minimum size for a lateral root to be detected as a branching root is the parent roots radius multiplied with 0.2 mm. The topological and morphological information are exported as CSV and the processed segments with the calculated skeleton is saved as PNG (Figure 2.4). The feature extraction is started from the “RhizoVision Explorer” GUI. Further details and background information are explained in (39).

### Root analysis

As last step in addition to the feature extraction, the two-dimensional root length density (RLD) is calculated from the total root length and the window size of the image in the unit of cm cm^−2^. Furthermore, the number of root tips and branch points, the total root length, the branching frequency, the network and surface area, the diameter, the perimeter and the volume are extracted from the “RhizoVision Explorer” output CSV and applied to spatio-temporal analysis of the root system (Figure 3).

**Fig. 3.**
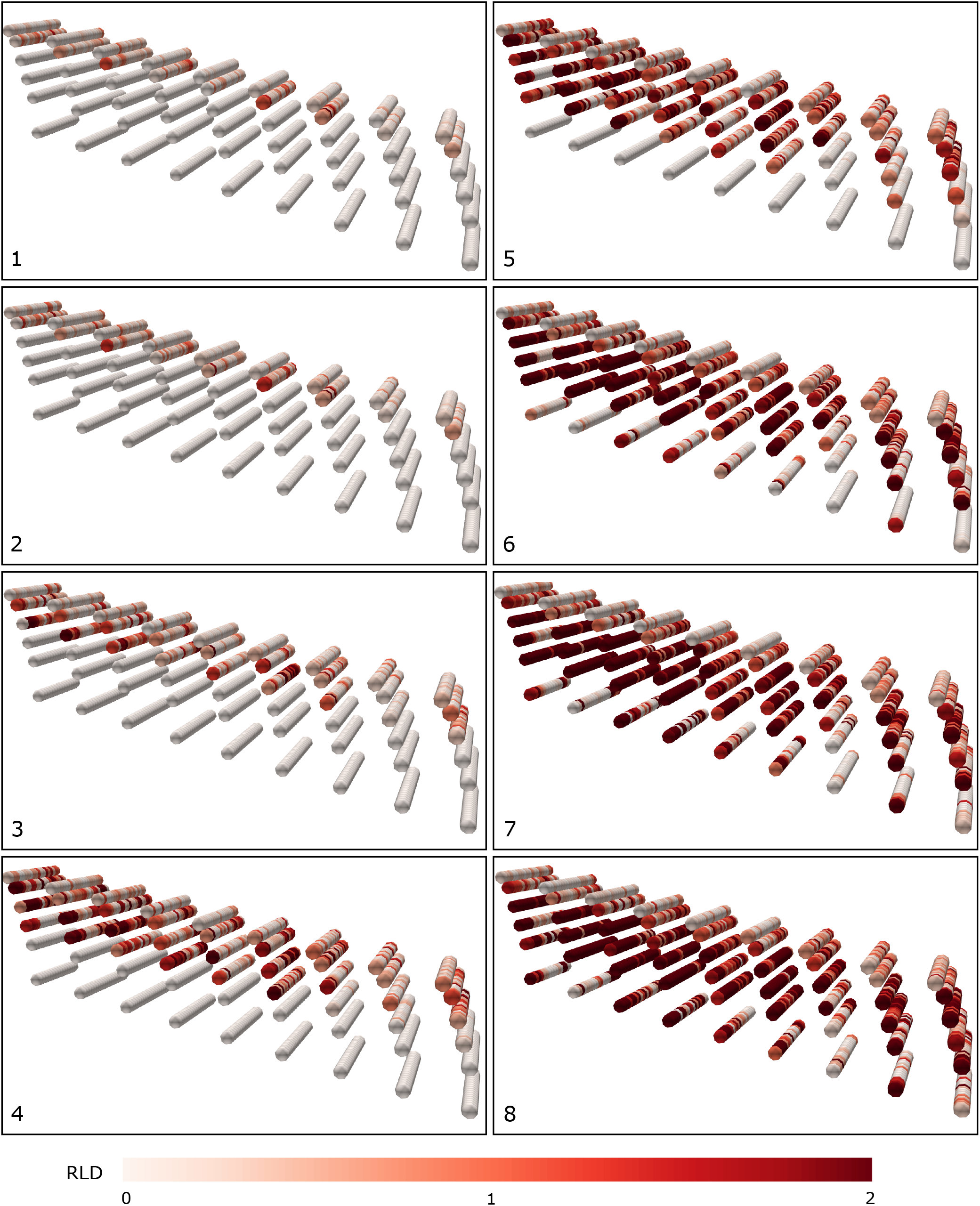
3D spatio-temporal distribution of RLD measured in all tubes at one minirhizotron. Distances between tubes are not to scale. (here: RLT in 2018). 1-8 represent the time steps.

### Evaluation

The F_1_-score (eqn. 1) is a measure commonly used to evaluate neural network models (27).

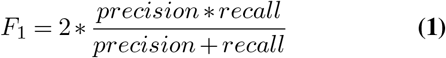

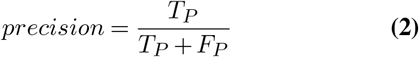

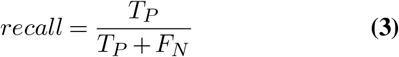

where *T_P_* are the *true positive*, *F_P_* the *false positive* and *F_N_* represent the *false negative* pixels recognized as root segment. The *F_1_* -score was calculated during the interactive training. Positive and negative values are the amount of pixels annotated in the interactive training as root and not-root.

The outcome of the automated root annotation was compared to the manual annotation by means of the pearson correlation coefficient, both on the data set as a whole as well as on individual growing seasons and measurement dates for the seasons 2017 and 2018. For the same seasons we calculated the mean root length per image at each measurement date and used a Welch two-sample t-test to assess whether the differences between manual and automated annotation of the total root length (ΔRL) were statistically significant. Furthermore the normalized root mean squared error (NRMSE) was calculated according to eqn. (4).

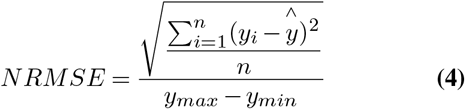

where *n = sample size, y_i_* is the *i^th^ observation of y* and 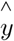 is the *predict y value*.

The manual per-image annotation with “Rootfly” of 2015/16 data is no longer available. However, the images and mean RLD values per tube are available and therefore were used for comparison. Based on this the RLD resulting from automated and manual analysis methods was calculated for every minirhizotron tube and measurement date (Figure 3) and compared as a proxy for a common root measurement parameter (40). In this analysis, all growing periods 2015/16, 2017 and 2018 were included.

### Statistics, data processing and visualization

Python 3.8 with *Pandas 1.0.5*, *Numpy 1.18.5*, *Matplotlib 3.2.2*, *Pillow 8.2.0* and *SciPy 1.5.0* have been used for statistics, data processing and visualization.

## Results

### Neural Network model validation

The *F_1_* for both neural network models trained for each camera-system is high. The F_1_ for the Bartz-system is 0.78 and 0.81 for VSI-system model.

### Comparison of automated and manual annotation

Generally, the Pearson-correlation of the total root length between manual annotation and automated analysis pipeline is high (0.7<R<0.9). Considering 58,360 images used for comparison, the overall correlation is *R* = 0.81.

By looking more into detail at each measurement date in 2017 and 2018, the correlation between manual annotation and the workflow for automated root segmentation is mostly high. Generally, the highest accordance and continuity between manual annotation and automated analysis pipeline was found in the results of the images taken in 2017. The correlations performed with the images of 2018 vary depending on measurement date and facility. Considering the measurements of 2017 separately, the correlation was performed with 16,599 images taken at R_UT_ and 20,388 images taken at R_LT_. For the dataset, analyzed from the images taken in the growing period 2017, the correlation is high to very high (*R* = 0.77 - 0.94) for every measurement date except the first measurement date at R_LT_ (*R* = 0.57) (Figure 7). For the correlation of the data, obtained from manual annotation and automated analysis of the images taken in 2018, 11,314 images taken at R_UT_ and 10,069 images taken at R_LT_ were analyzed. In 2018 the correlation is still mostly high at R_UT_ for the measurement dates at the mid section of the growing period (*R* = 0.7 - 0.74), while the correlation at very early and very late measurement dates is only medium (0.5<R<0.7). Regarding the measurements at R_LT_, the correlation is lower (*R* = 0.2 - 0.72) with increasing tendencies from first to last measurement date (Figure 8). Generally, the correlation shows an increasing trend towards later measurement dates.

Regarding the RLD values from 2015/16, one specific difference between manual and automated analysis is visible. Until the 14*^th^* measurement date the RLD is continuously increasing and then stagnating in the 2015/16 data resulting from manual annotation. The RLD from the automated analysis follows the same trend but decreases from 14^*th*^ measurement date continuously. Beyond this, the RLD curves of both methods are very consistent (Figure 4). In 2017 datasets, only negligible differences between manual and automated analysis method are recognizable, except for the first measurement date at R_LT_ (Figure 5b,Figure 9b) and first two dates and a small peak at the fourth measurement at R_UT_ (Figure 5d). The data originating from 2018 show more irregularities between both methods (Figure 5e-h,Figure 9e-h). Especially a decreasing trend in late state of growing period (R_UT_) (Figure 9f) and that although the trend was similar, the absolute values of the RLD are systematically larger in the automated than in the manual annotation at R_LT_ (Figure 5h,Figure 9h). This parallel shift is the highest divergence between the methods which is also indicated by the NRMSE and the differences in the mean of total root length (Table 2).

**Fig. 4.**
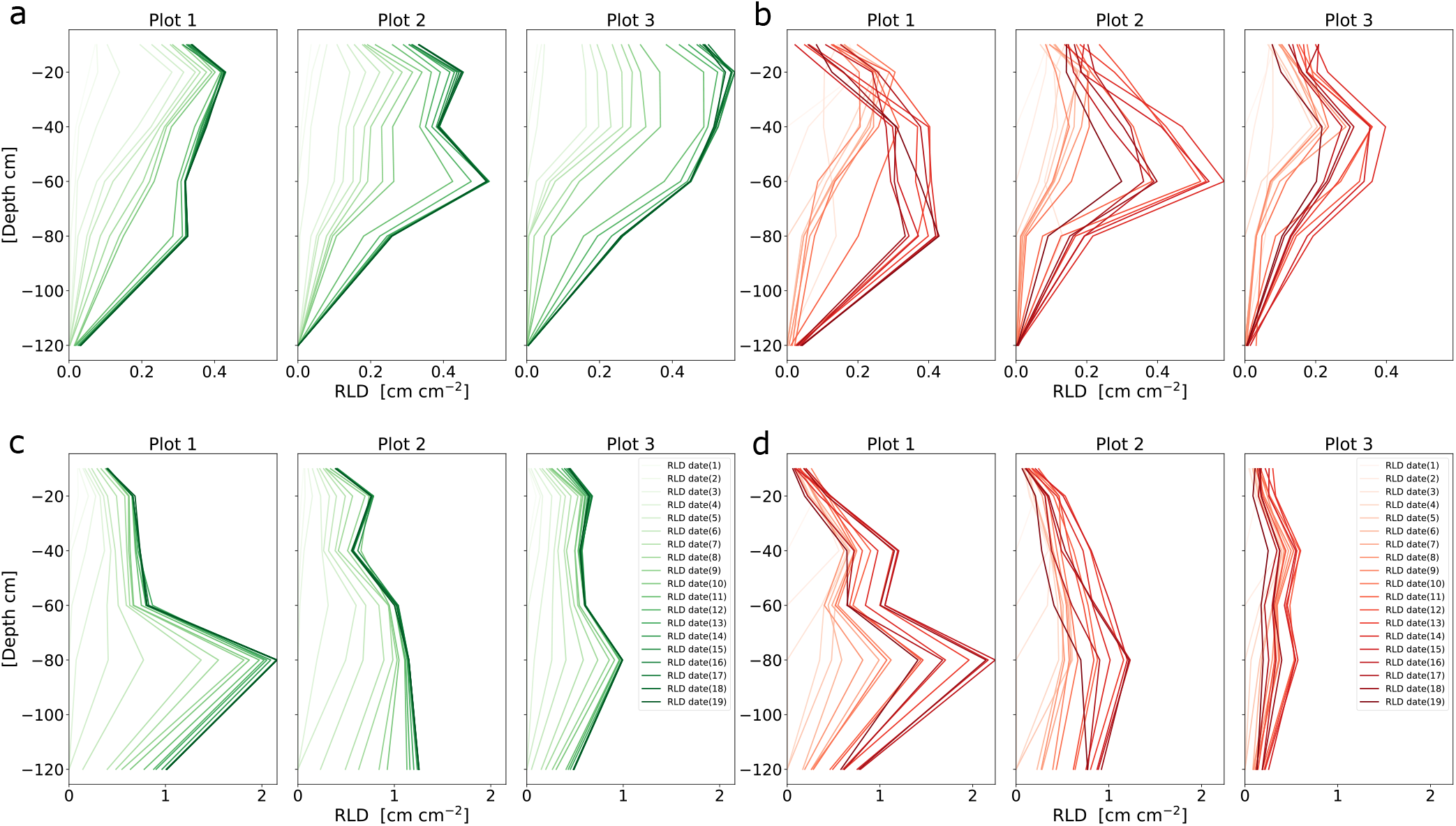
Comparison RLD of the data obtained from images originating from two minirhizotrons in the growing season 2015/16, separated by plots grown with different cultivars. The images were analyzed by hand (left (green): manual) and by the automated analysis pipeline (right (red): automated). a) R_UT_ manual, b) R_UT_ automated, c) R_LT_ manual,d) R_LT_ automated

**Fig. 5.**
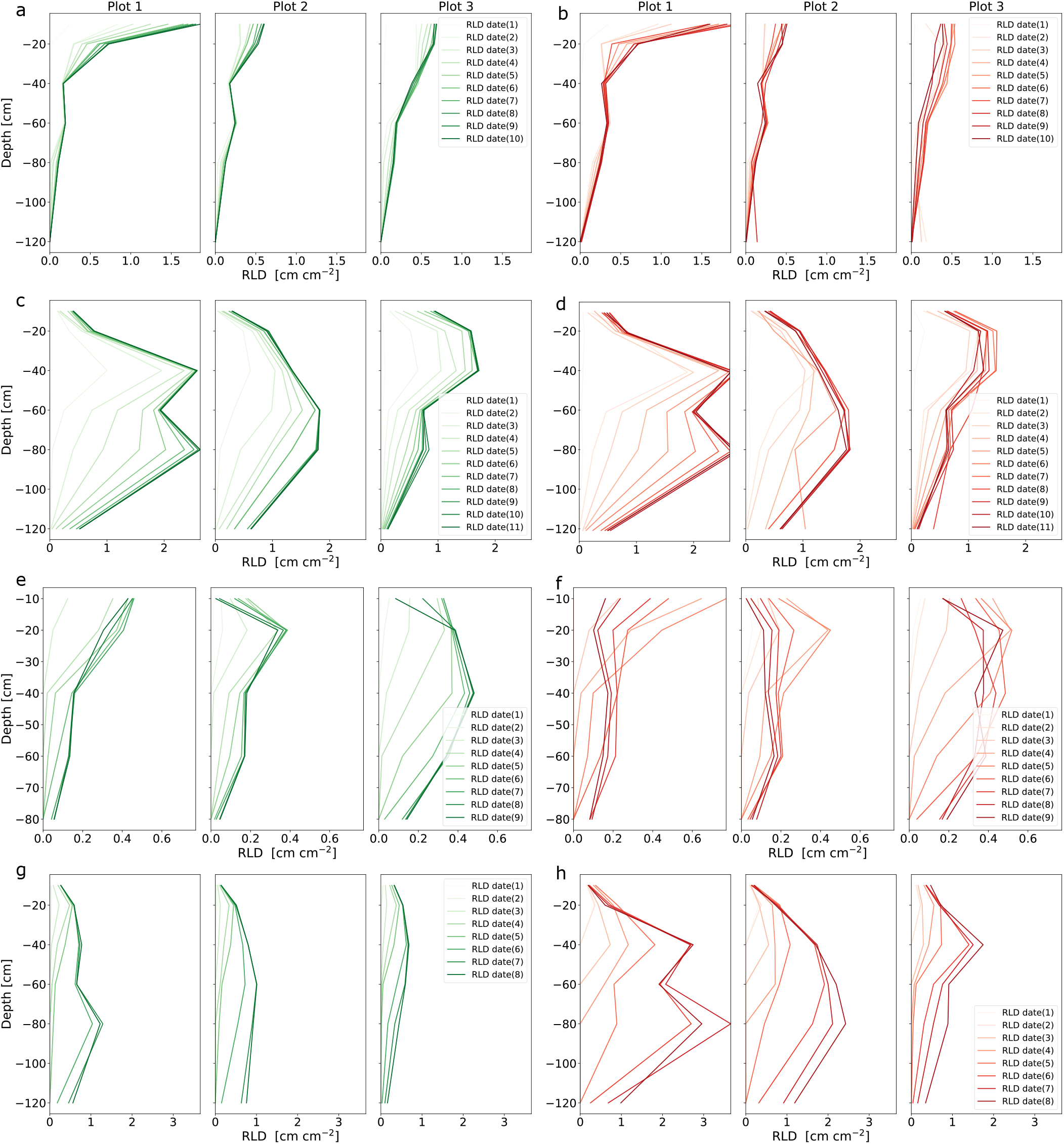
Comparison RLD of the data obtained from images originating from two minirhizotrons in the growing seasons 2017 and 2018, separated by plots grown with different cultivars. The images were analyzed by hand (left (green): manual) and by the automated analysis pipeline (right (red): automated). 2017: a) R_UT_ manual, b) R_UT_ automated, c) R_LT_ manual,d) R_LT_ automated; 2018: e) R_UT_ manual, f) R_UT_ automated, g) R_LT_ manual, h) R_LT_ automated.

**Table 2.** Overview of the statistical comparison of automated and manual annotation. ΔRL is the difference between the mean total root length (mm) obtained from manual and automated analysis methods, and a Welsch two sample t-test shows whether differences are significant (* = p<0.01).

|  |  | 2017 |  | 2018 |  |
| --- | --- | --- | --- | --- | --- |
| sampling # |  | R<sub>UT</sub> | R<sub>LT</sub> | R<sub>UT</sub> | R<sub>LT</sub> |
| 1 | $\Delta RL$ | 4.25* | 7.27* | 0.35* | 2.49* |
|  | NRMSE | 0.048 | 0.204 | 0.070 | 0.143 |
|  | R | 0.92 | 0.53 | 0.38 | 0.2 |
| 2 | $\Delta RL$ | 0.89 | 13.05* | 0.31 | 2.56* |
|  | NRMSE | 0.071 | 0.105 | 0.066 | 0.134 |
|  | R | 0.78 | 0.83 | 0.61 | 0.36 |
| 3 | $\Delta RL$ | 0.95 | 0.54 | 0.8 | 4.88* |
|  | NRMSE | 0.057 | 0.052 | 0.078 | 0.165 |
|  | R | 0.83 | 0.92 | 0.74 | 0.45 |
| 4 | $\Delta RL$ | 2.94* | 0.65 | 2.34 | 8.96* |
|  | NRMSE | 0.051 | 0.055 | 0.092 | 0.209 |
|  | R | 0.88 | 0.9 | 0.74 | 0.54 |
| 5 | $\Delta RL$ | 1.36 | 0.92 | 2.54* | 14.36* |
|  | NRMSE | 0.041 | 0.072 | 0.097 | 0.292 |
|  | R | 0.9 | 0.87 | 0.75 | 0.57 |
| 6 | $\Delta RL$ | 1.35 | 1.7 | 0.67 | 22.8* |
|  | NRMSE | 0.044 | 0.065 | 0.091 | 0.354 |
|  | R | 0.9 | 0.84 | 0.74 | 0.72 |
| 7 | $\Delta RL$ | -1.46 | 1.8 | -0.88 | 24.75* |
|  | NRMSE | 0.046 | 0.058 | 0.079 | 0.196 |
|  | R | 0.88 | 0.93 | 0.7 | 0.72 |
| 8 | $\Delta RL$ | 0.11 | 0.55 | -1.76* | 26.61* |
|  | NRMSE | 0.045 | 0.073 | 0.093 | 0.340 |
|  | R | 0.86 | 0.89 | 0.7 | 0.68 |
| 9 | $\Delta RL$ | -1.41 | -0.97 | -1.53 | |
|  | NRMSE | 0.039 | 0.057 | 0.103 |  |
|  | R | 0.88 | 0.94 | 0.63 |  |
| 10 | $\Delta RL$ | -2.44 | -2.89 | | |
|  | NRMSE | 0.039 | 0.047 |  |  |
|  | R | 0.88 | 0.92 |  |  |
| 11 | $\Delta RL$ | | 0.65 | | |
|  | NRMSE |  | 0.065 |  |  |
|  | R |  | 0.92 |  |  |

**Table 3.** Detailed overview of the images taken at the growing season 2015/16, 2017 and 2018.

| 2015/16 |  |  |  | 2017 |  | 2018 |  |
| --- | --- | --- | --- | --- | --- | --- | --- |
| no. | facility | date | images | date | images | date | images |
| 1 | R<sub>UT</sub> | 16/11/15 | 720 | 08/06/17 | 480 | 23/05/18 | 440 |
|  | R<sub>LT</sub> | 16/11/15 | 720 | 08/06/17 | 584 | 23/05/18 | 720 |
| 2 | R<sub>UT</sub> | 26/11/15 | 1,080 | 29/06/17 | 1,800 | 20/05/18 | 480 |
|  | R<sub>LT</sub> | 26/11/15 | 1,079 | 22/06/17 | 1,800 | 30/05/18 | 720 |
| 3 | R<sub>UT</sub> | 17/12/15 | 1,800 | 06/07/17 | 1,800 | 07/06/18 | 960 |
|  | R<sub>LT</sub> | 17/12/15 | 1,439 | 29/06/17 | 2,160 | 07/06/18 | 1,075 |
| 4 | R<sub>UT</sub> | 02/02/16 | 1,520 | 13/07/17 | 1,800 | 18/06/18 | 1,280 |
|  | R<sub>LT</sub> | 21/01/16 | 1,800 | 06/07/17 | 2,160 | 18/06/18 | 1,439 |
| 5 | R<sub>UT</sub> | 12/02/16 | 1,800 | 20/07/17 | 1,800 | 26/06/18 | 1,400 |
|  | R<sub>LT</sub> | 12/02/16 | 1,800 | 13/07/17 | 2,160 | 26/06/18 | 1,800 |
| 6 | R<sub>UT</sub> | 26/02/16 | 1,800 | 27/07/17 | 1,200 | 07/07/18 | 1,638 |
|  | R<sub>LT</sub> | 26/02/16 | 2,160 | 20/07/13 | 2,160 | 18/07/18 | 2,156 |
| 7 | R<sub>UT</sub> | 14/03/16 | 1,800 | 02/08/17 | 1,840 | 18/07/18 | 1,760 |
|  | R<sub>LT</sub> | 14/03/16 | 2,160 | 27/07/17 | 1,430 | 01/08/18 | 2,159 |
| 8 | R<sub>UT</sub> | 26/03/16 | 1,840 | 10/08/17 | 1,959 | 01/08/18 | 1,680 |
|  | R<sub>LT</sub> | 24/03/16 | 2,160 | 02/08/17 | 2,159 | 23/08/18 | 2,159 |
| 9 | R<sub>UT</sub> | 07/04/16 | 2,160 | 23/08/17 | 2,120 | 16/08/18 | 1,676 |
|  | R<sub>LT</sub> | 07/04/16 | 2,160 | 10/08/17 | 2,160 | - | - |
| 10 | R<sub>UT</sub> | 13/04/16 | 2,160 | 12/09/17 | 1,800 | - | - |
|  | R<sub>LT</sub> | 13/04/16 | 2,160 | 24/08/17 | 2,159 | - | - |
| 11 | R<sub>UT</sub> | 29/04/16 | 2,160 | - | - | - | - |
|  | R<sub>LT</sub> | 29/04/16 | 2,160 | 12/09/17 | 2,150 | - | - |
| 12 | R<sub>UT</sub> | 06/05/16 | 2,160 | - | - | - | - |
|  | R<sub>LT</sub> | 06/05/16 | 2,160 | - | - | - | - |
| 13 | R<sub>UT</sub> | 13/05/16 | 2,160 | - | - | - | - |
|  | R<sub>LT</sub> | 13/05/16 | 2,160 | - | - | - | - |
| 14 | R<sub>UT</sub> | 20/05/16 | 2,160 | - | - | - | - |
|  | R<sub>LT</sub> | 20/05/16 | 2,160 | - | - | - | - |
| 15 | R<sub>UT</sub> | 27/05/16 | 2,160 | - | - | - | - |
|  | R<sub>LT</sub> | 27/05/16 | 2,159 | - | - | - | - |
| 16 | R<sub>UT</sub> | 03/06/16 | 2,160 | - | - | - | - |
|  | R<sub>LT</sub> | 03/06/16 | 2,159 | - | - | - | - |
| 17 | R<sub>UT</sub> | 09/06/16 | 2,160 | - | - | - | - |
|  | R<sub>LT</sub> | 09/06/16 | 2,160 | - | - | - | - |
| 18 | R<sub>UT</sub> | 16/06/16 | 2,155 | - | - | - | - |
|  | R<sub>LT</sub> | 16/06/16 | 2,160 | - | - | - | - |
| 19 | R<sub>UT</sub> | 23/06/16 | 2,149 | - | - | - | - |
|  | R<sub>LT</sub> | 23/06/16 | 2,156 | - | - | - | - |

The comparison between two human annotators and each annotator and the automated analysis pipeline separately shows that the correlation between the person 1 and the pipeline is *R* = 0.92 and the correlation between person 2 and the pipeline is *R* = 0.79. The correlation between both persons is the lowest (*R* = 0.73).

### Time evaluation

The time required to train the neural network model mostly depends on the amount of images included in the training dataset. Approximately 65 % of the time needed is used for training of the deep neural network. The annotation takes 40 % of the time, based on a mean of 200 annotated images h^−1^. The mean time needed by the network for the training of a dataset of 1,500 images, was approximately 5h, excluding the real-time training during the annotation. This is approximately 25 % of the entire processing time. Segmentation took around 27 % of the total time. With 4 Nvidia GeForce RTX 2080 Ti GPUs and a batch size of 12 the segmentation took around 0.7 s. Converting the segments to binary and the final feature extraction took around 8 %of the time (Figure 6).

**Fig. 6.**
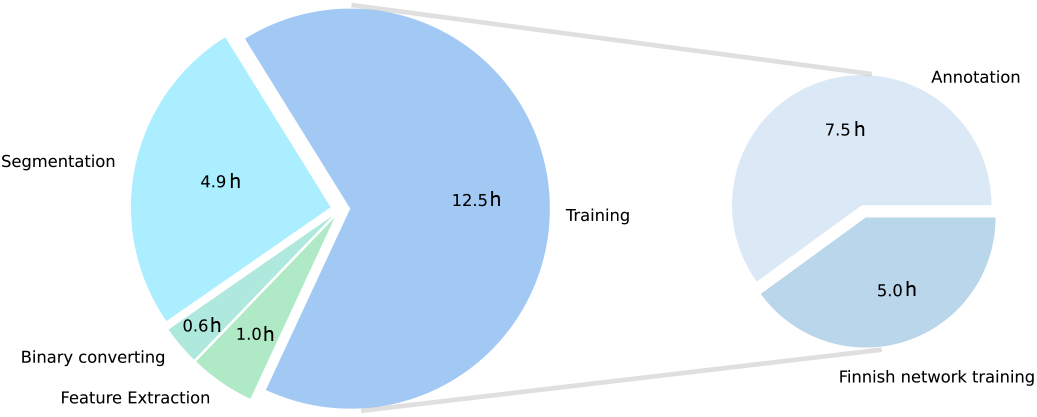
Time requirements to run the automated analysis pipeline for a sample of 25,000 images. Left: All sub-processes together. Right: share of the neural network training, which is only required when no suitable model is available.

## Discussion

### Availability and feasibility

The availability is the parameter for how easily accessible all components of the automated pipeline are for everyone. The feasibility defines how easy the proposed pipeline and with that the required software can be operated. The equipment needed to apply the new workflow requires a computer with a powerful GPU, or alternatively a basic computer, an additional server with powerful GPUs and a network-connection between both. Furthermore, the software packages of “RootPainter” and “RhizoVision Explorer” are needed and the conversion and analysis script are required. All this is open-source available (16, 29). Once the model is trained, the human interaction needed to apply the pipeline is reduced to a few “clicks”. With a suitable model available, the user has to interact actively three times with the automated pipeline, (*1*.) to start the segmentation, (*2*.) to convert the segments to binary and (*3*.) to start the feature extraction. No deeper knowledge in computer science is needed, because all intermediate steps are available within a GUI. However, the first implementation of the “RootPainter” environment at the server part of the setup requires basic knowledge in server administration or support. In contrast to manual or semi-automated operated root analysis programs, like different tools based on “ImageJ”, “DART”, “GiA Roots”, “SmartRoot”, “EZ-Rhizo” or “Rootfly”, the expenses in man-power, time, knowledge and experiences required to apply the automated workflow, are much lower. This is granted due to the very small interactions needed for the automated analysis pipeline (13, 28, 41–44).

### Accuracy and comparability

The accuracy evaluates the automated analysis pipeline in terms of reliability and exactness of the generated data. Comparability is given, if the results of the automated analysis pipeline can be compared to the outcome of previously evaluated data of the same kind, like the manual annotation performed with “Rootfly”. The most important characteristic of the automation of plant data analysis is the reliability of the generated datasets. Therefore, the accuracy of the observed root traits has to be as close to the ground truth as possible (5). In our study we used the manual annotation of the roots as comparison. The manual annotation was performed by different persons and over a long time period. Consequently, a certain subjectivity was included in this process.

Our data indicate that our pipeline provides a high tolerance towards different environmental conditions, explicitly different soil conditions. This is shown by the over-all correlation difference between the minirhizotrons in silty gravel soil and in gravely loam soil that is only 0.04. Regarding the corresponding measurement dates at R_UT_ and R_LT_ in 2017, there is only one date where the correlation difference of automated pipeline and manual annotation is larger than 0.1 between R_UT_ and R_LT_ (Figure 7). This gets even more visible by comparing the average difference in mean between the manual annotation and the automated workflow. Low difference in sample means, paired with the high correlation, shows a robust stability of this model against different soil-, respectively background-conditions. Generally, the results for 2017 data analyzed automatically and manually are impressively close to each other, indicating a general great fit of the models used for images originating from 2017. The impressive consistency of the automated analysis results becomes especially visible regarding the RLD plots plottet from 2015/16 and 2017 data (Figure 4, Figure 5). The decrease in 2015/16 RLD profiles that is not monitored in the manual annotation data, originated from the root senescence. The senescence could be better evaluated by the neural network than by the human annotator. In manual annotation the slight, gradual discoloration of the roots visually revealing the senescence is easy to miss. Furthermore, it is a complicated work step in “Rootfly” to eliminate already annotated roots at the right point in the timeline. Taking this into consideration, the results of the method comparison for 2015/16 and 2017 data shows impressive results, regarding accuracy and comparability of the automated analysis pipeline.

**Fig. 7.**
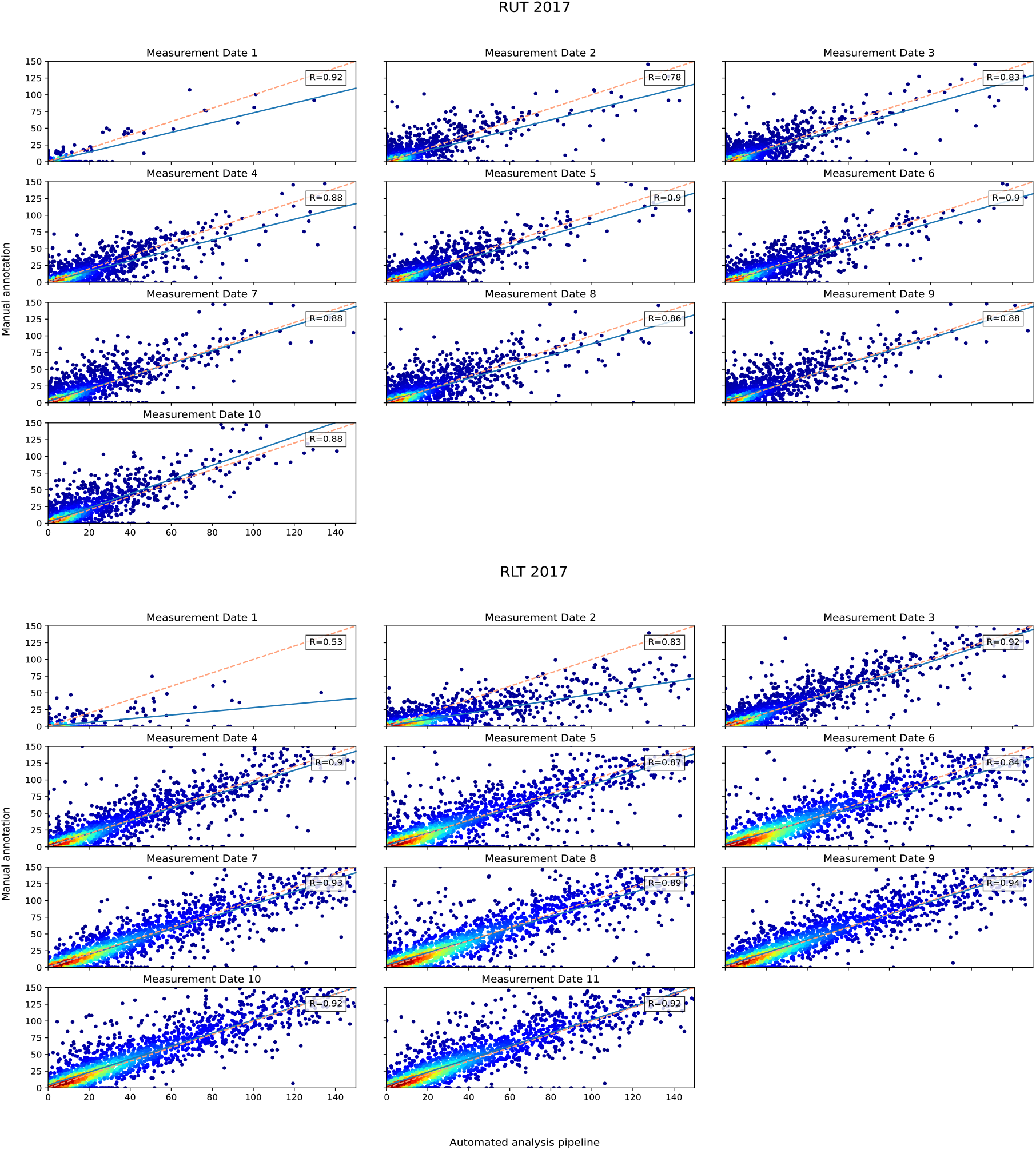
Correlation of automated and manual analyzed root length, obtained from 2017. Each measurement date is considered separately for R_UT_ and R_LT_. The color represents the density.

**Fig. 8.**
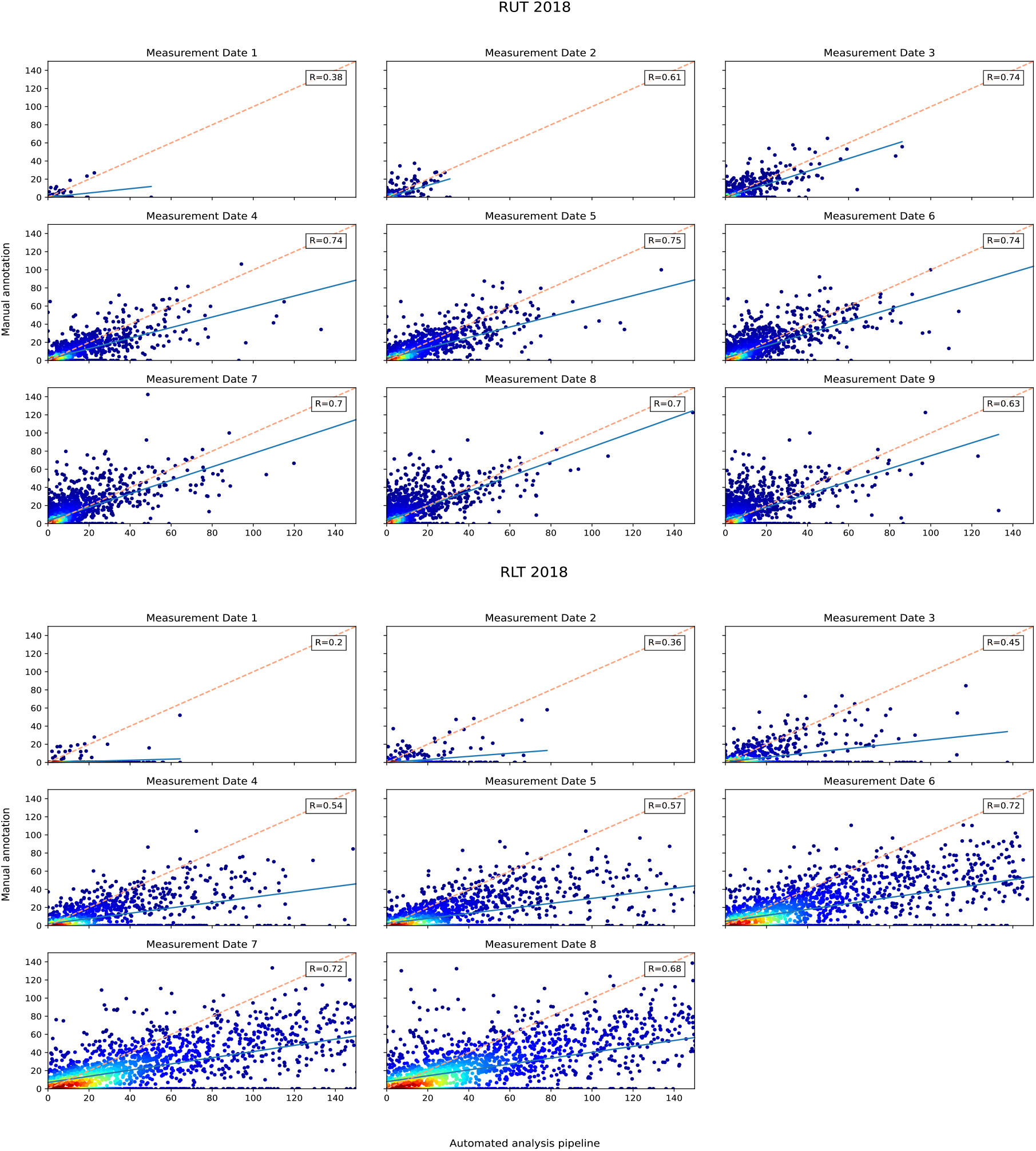
Correlation of automated and manual analyzed root length, obtained from 2018. Each measurement date is considered separately for R_UT_ and R_LT_. The color represents the density.

**Fig. 9.**
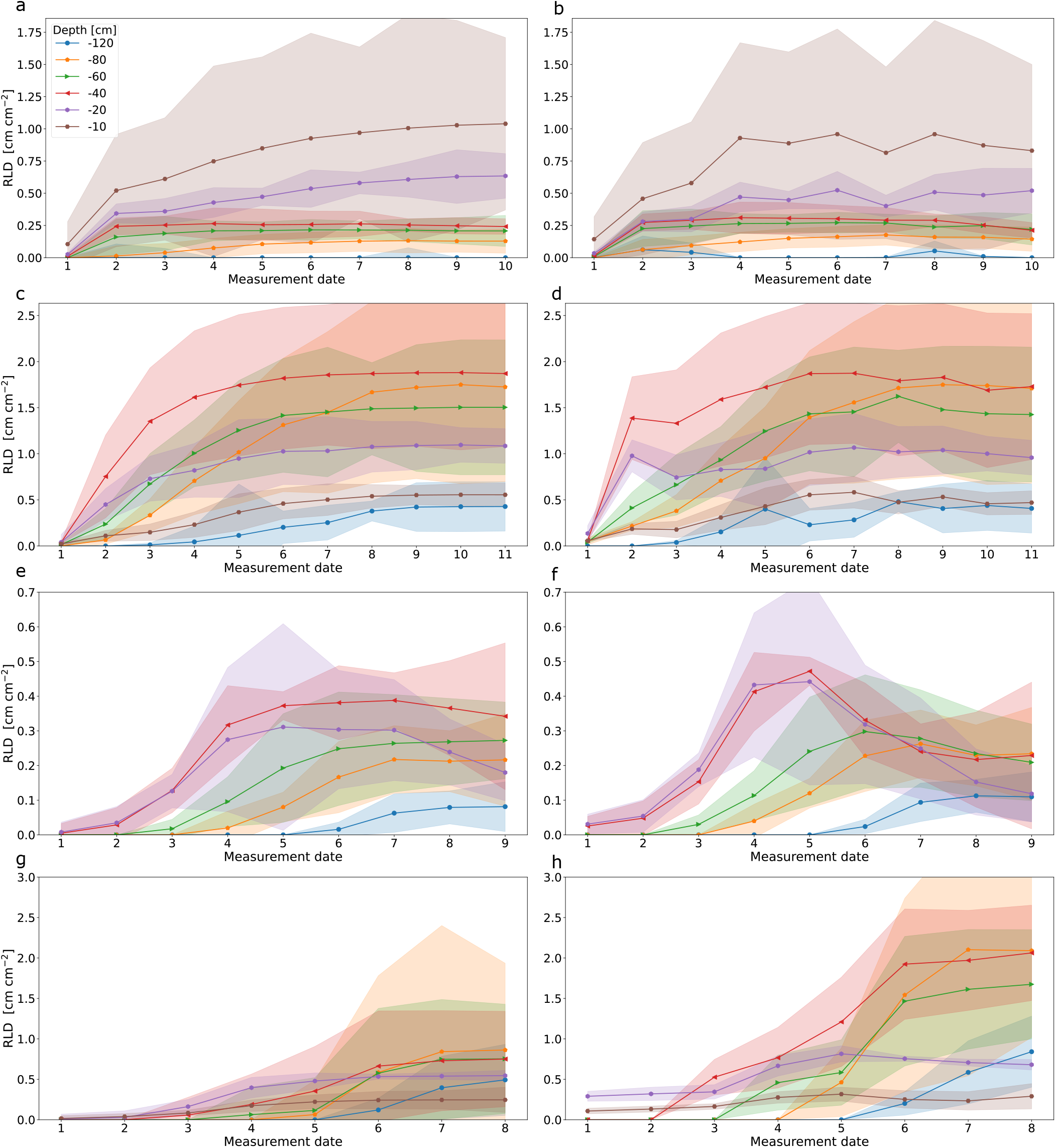
Comparison of root arrival curves of the data obtained from images originating from two minirhizotrons in the growing seasons 2017 and 2018. The images were analyzed by hand (left: manual) and by the automated analysis pipeline (right: automated). 2017: a) R_UT_ manual, b) R_UT_ automated, c) R_LT_ manual,d) R_LT_ automated; 2018: e) R_UT_ manual, f) R_UT_ automated, g) R_LT_ manual, h) R_LT_ automated. The shaded area represents the standard deviation.

Regarding the results obtained from the method comparison of 2018 data, this conclusion cannot be transferred in the same way.

The errors in our results of 2018 data show up especially at R_LT_ data, but are also visual at R_UT_. The discrepancies mainly originate from three facts. First, the visual inspection of the root segments revealed that old and not completely degraded root fragments partially caused errors at the very beginning of the vegetation period. Consequently, this occurs especially on images taken early in the growing season. The trained CNN sometimes segmented a small part of the residuals of old roots, especially at the first measurements of each growing period. In later measurement dates this overestimation vanished. This was more likely to occur in 2018 data in contrast to 2017 data, because in 2017 the previous crop was winter wheat which was harvested in summer. Between harvest in 2016 and new sowing of maize in 2017 the degradation could proceed further than between harvest in 2017 and sowing in 2018. A second explanation for the observed differences in root length can be found in the process of manual annotation. The manual annotation itself requires a certain level of expertise in root phenotyping. This expertise is gained with a lot of personal experiences (8, 13). Therefore, it can be hypothesized that there is also a significant influence of subjectivity in human annotation. Over the years, different persons annotated the root datasets. Hence, the impact of differences resulting from varying manual annotation strategies might influence the results more than the differences between manual and automated analysis. The direct comparison between two annotators showed a higher difference between the persons annotating, than between the automated analysis pipeline and each human annotator. Consequently, we concluded that the human effect on manual annotation is higher than the impact of a mistake done by the automated workflow. This is substantiated by comparing the manual annotations of the neural network training images to the results of the neural network applied to the same images. Although the fitting of the neural network model to the annotation done in “RootPainter” to train the neural network was good, as the *F1*-score indicated, the manual annotation previously done in “Rootfly” differ in 2018, for the images obtained at R_LT_, significantly. Regarding especially the ΔRL it can be seen, that the differences in mean between manual annotation and automated analysis pipeline in 2017 are very low −0.4 mm (R_UT_), 1.55 mm (R_LT_) and 0.84 mm for R_UT_ in 2018, while ΔRL = 19.34 mm for the images obtained from R_LT_ in 2018. This also indicates that the divergence between automated and manual analysis most likely result from the subjective annotation behavior of different annotators. In our case, it is most likely that an at least slightly different scheme of root detection were applied systematically to the manual annotation of 2018, than to the training to the neural network. Partly this is rooted in the subjective perception of the annotators, however this cannot explain the divergences to the fullest extent. The third and most affecting reason for the worse fit, is the annotation style generally used in 2018 data. The annotator only annotated the main roots and first order roots, while the automated analysis pipeline also detects all fine roots. The resulting higher values provided by the automated analysis are therefore more likely to fit with the real values in field than the manual annotation. This effect is stronger at R_LT_, because there are generally more roots, due to a better soil quality (6). These three influential factors distort the results of the manual annotation of images taken in 2018. Nevertheless, the pipeline provides reasonable results for these dataset and moreover outperforms the manual annotation especially, because it provides a holistic analysis by including all roots.

The automated analysis pipeline provides a level of objectivity, a human annotator cannot achieve. Therefore, it is highly probable that with the application of the automated pipeline associated minimization of the human influence will significantly improve objectivity and also accuracy of the minirhizotron image analysis.

### Speed and efficiency

The speed is the pure amount time the pipeline requires to analyses a certain amount of images. Efficiency is defined through the amount of time and labor needed to analyses a dataset in contrast to manual annotation. The time required to analyze root images by hand is enormous. The estimated time to analyze 100 cm^2^ of depicted soil is 1 - 1,5 h (16). This is consisted with the results of other studies, needing approximately 1 h for annotating 17-38 images manually (15). Intern evaluation reproduced the same results. To annotate 25,000 images, which is approximately the amount of images for a shorter growing season, the annotation time needed is 1000 - 1500 h. The time needed to process the same amount of images with the automated pipeline is approximately 19 h, including the training of the neural network. Without the training, the segmentation and feature extraction would only take around 6,5 h for all images. The resulting benefits in time saving are massive (Figure 6).

Generally, only around 1.2 % - 1.9 % of the time needed for manual annotation is needed by the automated workflow to process the data, including the training. Excluding the entire training process, the automated workflow requires only 0.4 % - 0.65 % of the time needed to annotate the same amount of images manually with e.g. “Rootfly”. Regarding the advantages of time saving, it further has to be taken into account that the time of interaction with the computer is decimated to almost zero, once the training is completed.

## Conclusion

We propose a new approach to analyze large amounts of 2D root image data. This became necessary with the big amount of data created in experimental field sites such as the minirhizotron facilities in Selhausen as well as others (45, 46). The automated analysis pipeline illustrated in this study, is a suitable solution to easily and accurately analyze minirhizotron images in significantly less time. To the best of our knowledge, we are the first study testing a deep learning and automated feature extraction combining high-throughput minirhizotron image analysis pipeline to this extend. The biggest advantage of the automated workflow is the massive saving in time. Precisely expressed, the required time is reduced by more than 98 % in contrast to manual annotation, while providing several root traits, including number of root tips, number of branch points, root length, branching frequency, network area, perimeter, volume, surface area and diameter on a spatio-temporal scale. The required root traits can be made available quickly which may speed up further analysis and applications of this type of data. In conclusion, the automated pipeline outperforms the manual annotation in time requirements and information density, while providing reliable data and feasibility for everyone. Tested with more than 131,000 minirhizotron images, including more than 58,000 images for detailed comparison, obtained from three growing seasons and different soil types, depths and cultures our results indicate a high general validity for the presented pipeline. Irregularities in the match of manual annotation and analysis pipeline can be essentially explained by rarely appearing false recognize of old decreasing roots and mainly by the influence of human subjectivity in annotation. Balanced training datasets and consequent annotation of the training data are the key to good results. If these facts are considered, the here presented and evaluated pipeline has the potential to be the new standard method for reliable high-throughput root phenotyping of (mini)rhizotron images.

## ACKNOWLEDGEMENTS

This work has partially been funded by the German Research Foundation under Germany’s Excellence Strategy, EXC-2070 - 390732324 - PhenoRob and by the German Federal Ministry of Education and Research (BMBF) in the framework of the funding initiative “Plant roots and soil ecosystems, significance of the rhizosphere for the bio-economy” (Rhizo4Bio), subproject CROP (ref. FKZ 031B0909A).

We thank Xavier Draye for the early access to the Manneback-cluster of the uCLou-vain. We thank Abraham Smith for the help with RootPainter and Larry York for the help with RhizoVision Explorer.

## Appendix

### Data availability

1. The datasets generated and/or analysed during the current study are available in this repository: https://hdl.handle.net/20.500.11952/butt.data.handle/00000051. If you only wish to use the data, please cite as: Bauer, Felix (2021): DATA: Combining deep learning and automated feature extraction to analyze minirhizotron images development and validation of a new pipeline (Dataset). Forschungszentrum Jülich, 10.34731/pbn7-8g89.
2. RootPainter (27) is available at: https://github.com/Abe404/root_painter
3. RhizoVision Explorer (29, 39) is available at: https://zenodo.org/record/4095629 and https://github.com/rootphenomicslab/RhizoVisionExplorer

